# Directly interfacing brain and deep networks exposes non-hierarchical visual processing

**DOI:** 10.1101/2021.06.28.450213

**Authors:** Nicholas J. Sexton, Bradley C. Love

## Abstract

One reason the mammalian visual system is viewed as hierarchical, such that successive stages of processing contain ever higher-level information, is because of functional correspondences with deep convolutional neural networks (DCNNs). However, these correspondences between brain and model activity involve shared, not task-relevant, variance. We propose a stricter test of correspondence: If a DCNN layer corresponds to a brain region, then replacing model activity with brain activity should successfully drive the DCNN’s object recognition decision. Using this approach on three datasets, we found all regions along the ventral visual stream best corresponded with later model layers, indicating all stages of processing contained higher-level information about object category. Time course analyses suggest long-range recurrent connections transmit object class information from late to early visual areas.

## Main Text

Despite some shortcomings (*1,2*), deep convolutional neural networks (DCNNs) have emerged as the best candidate models for the mammalian visual system. These models take photographic stimuli as input and, after traversing multiple layers consisting of millions of connection weights, output a class or category label. Weights are trained on large datasets consisting of natural images and corresponding labels.

The deep learning revolution in neuroscience began when layers of DCNNs were related to regions along the ventral visual stream in an early-to-early and late-to-late pattern of correspondence between brain regions and model layers (3–5, fig. 1A). This correspondence supported the view that the ventral stream is a hierarchy in which ever more complex features and higher-level information are encoded as one moves from early visual areas like V1 or V4 to inferotemporal (IT) cortex (*6–8*).

**Figure 1:**
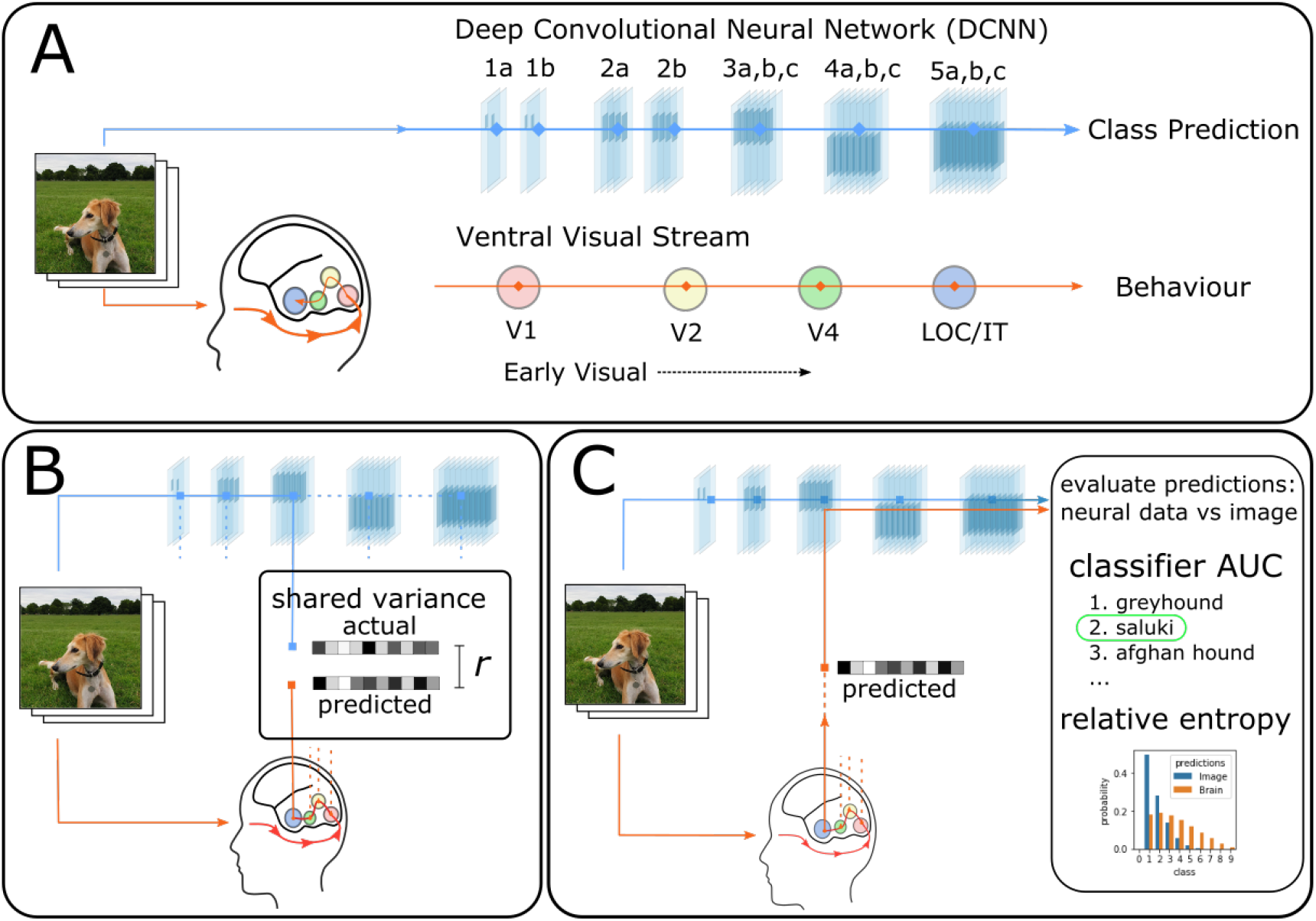
Deep Convolutional Neural Networks (DCNNs) trained on large naturalistic image datasets (*24*) have emerged as leading models of the mammalian ventral visual stream. (**A**) Typically, processing in DCNNs is hierarchical starting with the stimulus and proceeding across successive layers as higher-level information is extracted, culminating in predicting the class label (*13*). Numerous analyses (*3–5*) based on shared variance suggest the brain follows related principles with an early-to-early, late-to-late pattern of correspondence between the ventral visual stream and DCNN layers. (**B**) These shared-variance correspondences are evaluated locally, typically involving one brain region and one model layer, with no recourse to behaviour (i.e., the object recognition decision). (**C**) We propose a stronger test of correspondence based on task-relevant variance. If a model layer and brain region correspond, then model activity replaced with brain activity should drive the DCNN to an appropriate output (i.e., decision). The quality of correspondence is evaluated by comparing DCNN performance when driven by a stimulus image vs. interfaced with brain activity.

However, these correspondences between brain and model activity were based on total shared variance as opposed to task-relevant variance (fig. 1B). Much of cortex-wide neural variance does not relate to the task of interest (*9*) and may co-vary with but not drive behaviour. Correspondences established by correlation alone do not necessitate that model layers and brain regions play the same functional role in the overall computation.

We propose a stronger test for evaluating how brain-like a model is. If, as is frequently claimed (*3–5*), a specific layer in a DCNN corresponds to a brain region, then it should be possible to substitute the activations on that layer with the corresponding brain activity and drive the DCNN to an appropriate output (cf. 10–12, fig 1C). For example, if we take V4 activity from a monkey viewing an image of a car and interface that brain activity with an intermediate DCNN layer hypothesised to correspond to V4, then the DCNN should respond “car” absent any image input. How well the DCNN performs when directly interfaced (through a simple linear mapping (see SI materials & methods), with the brain provides a strong test of how well the interfaced brain region corresponds to that layer of the DCNN.

We interfaced a pretrained DCNN (*13*) with data from two human brain imaging studies (*14,15*) and a Macaque monkey study (*16*). All three studies involved viewing complex images. For a chosen model layer and brain region, we calculated a linear mapping from brain to model activity by presenting the same images to the model for which we had neural recordings (fig. 1C). This simple linear mapping is a translation between brain and model activity. We evaluated the quality of this translation by considering held-out images and brain data that were not used in calculating the linear mapping (SI materials & methods).

Strikingly, for the two fMRI studies (figs 2A, 2B), the DCNN was most accurate at classifying novel images when brain activity across regions (both early and late along the ventral stream) was interfaced with later model layers. In contrast to previous analyses that focused on total variance, we did not find the early-to-early and late-to-late pattern of correspondence. Even primary visual cortex, V1, best drove the DCNN when interfaced with an advanced layer. For comparison, classifiers commonly used to decode information from fMRI data through multivariate pattern analaysis (MVPA) were at chance levels (fig. S2), which highlights the useful constraints captured in the pretrained DCNN. After training on a million naturalistic images, the DCNN developed representations that paralleled those of the ventral stream, which made decoding object class possible by way of a linear mapping from brain activity to an advanced DCNN layer.

**Figure 2:**
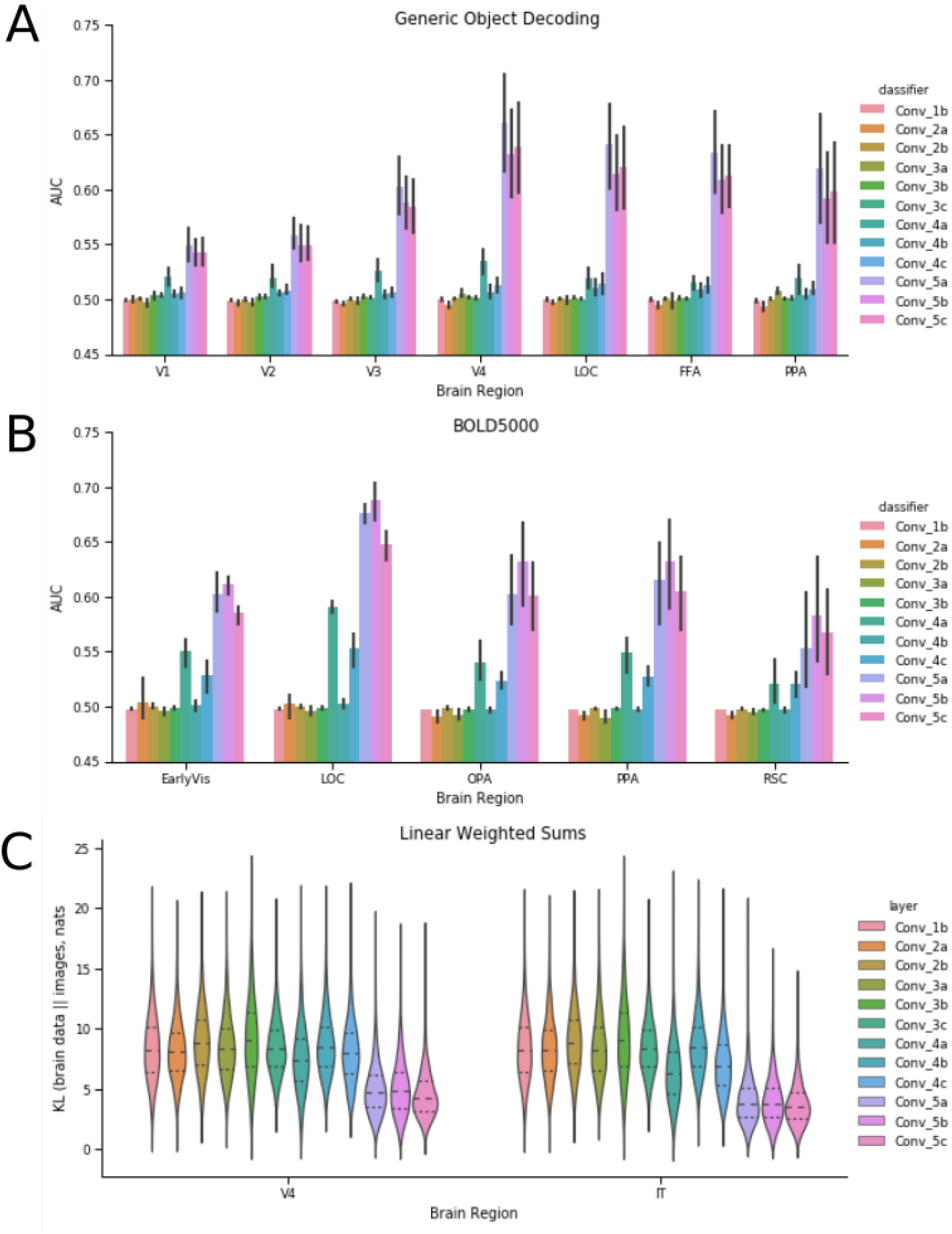
Results from interfacing neural data with a Deep Convolutional Neural Network (DCNN). Using the method shown in fig. 1C, brain activity is directly inputted to a model layer to assess correspondence between a brain region and model layer. (**A**) For this human fMRI study (*15*), all brain areas drive DCNN object recognition performance to above chance levels. Performance is best for all brain areas when interfaced with later model layers. (**B**) The same pattern of results is found for a second human fMRI study (*14*). (**C**) In a third study, KL divergence is used (see main text and SI) to measure the degree of correspondence for when the DCNN is driven by image input vs. multi-unit recordings from macaque monkeys (*16*). For KL divergence, lower values indicate better correspondence. Once again, all regions best correspond to later network areas. These three analyses indicate that higher-level visual information is present at all stages along the ventral visual stream.

The interpretation is that all brain regions contain advanced object recognition information, which conflicts with strict hierarchical views of the ventral visual stream.

To rule out any alternative explanation based on the indirect nature of fMRI recordings, we considered a third study consisting of direct multi-unit recording of spiking neurons implanted in the ventral visual stream of Macaque monkeys (*16*). These monkeys were shown images that did not readily align with the pretrained DCNN’s class labels, so we evaluated neural translation performance by comparing the outputs of the DCNN when its input was a study image vs. when a DCNN layer was driven by brain data elicited by the same image. For the distance measure, KL divergence, lower values imply a better translation between brain and model activity. As in the fMRI studies, both relatively early regions (i.e., V4) and late regions (i.e., IT) best translated to later DCNN layers (fig 2C).

Across three diverse studies, we found a remarkably consistent pattern that strongly diverged from previous analyses — both early and late regions along the ventral visual stream best corresponded (i.e., translated) to late model layers. It is not that previous analyses were poorly conducted (see fig S1 for a successful reanalysis of data (*16*) finding the early-to-early and late-to-late canonical pattern). Rather, our novel analyses focused on task-relevant analysis, i.e., variance that can drive behaviour, provided a different view of the system than standard analyses focused on shared variance. Integrating these two views suggests a non-hierarchical account of object recognition marked by long-range recurrence transmitting higher-level information to the earliest visual areas.

One way to reconcile the existing literature based on shared variance with our analyses based on task-relevant variance is to propose that long-range connections from IT transmit higher-level information to early visual areas. Even if most variance in lower-level visual areas is attributable to stimulus-driven, bottom-up activity, the majority of task-relevant information could be attributable to signals originating from IT (fig. 3).

**Figure 3:**
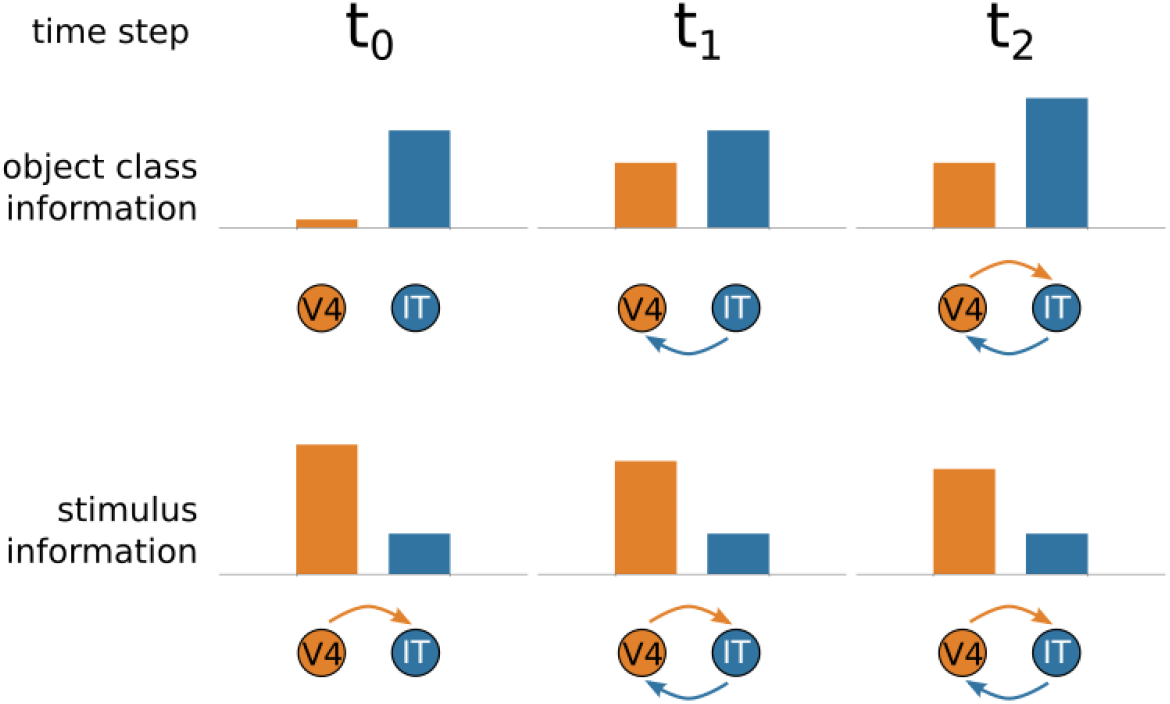
Hypothesised interactions between early (V4) and late (IT) regions along the ventral visual stream as processing unfolds. We hypothesise how stimulus and object-class information propagates between V4 and IT over time. At *t*_0_, the forward pass reaches IT from V4, with V4 activity reflecting low-level stimulus properties but little information about object class. At *t*_1_, object-class information from IT flows back to V4, increasing its task-relevant activity, which in turn influences IT at *t*_2_. Notice that later in processing, V4 reflects object class information, but most of its activity remains tied to bottom-up stimulus properties. These hypothesised interactions would reconcile our results (fig. 2) based on task-relevant information with previous results based on shared variance.

This view predicts specific patterns of Granger causality between early and late areas along the ventral visual stream. Do past values of one time series predict future values of the other? In terms of total spiking activity, lower-level areas should first cause activity in higher-level areas during the initial feed-forward pass in which stimulus-driven activity propagates along the ventral visual stream. Later in processing, the causality should become reciprocal as top-down connections from IT affect firing rates in lower-level areas, such as V4 (fig 3, bottom row). In contrast, Granger causality for task-relevant information should first be established from IT to V4 (i.e., the top-down signal) and only later in processing should recurrent activity lead to causality from V4 to IT (fig. 3, top row). In this fashion, all areas are effectively “late” after long-range recurrent connections transmit information from IT to early visual areas along the ventral stream though most variance for these areas would be dominated by lower-level (bottom-up) stimulus information.

We tested these predictions using the monkey multi-unit spiking data (*16*) that has the temporal resolution to support the analyses. Images were presented one after the other, each visible for 100ms, with a 100ms period between stimuli. Figure 4A shows the mean firing rates (10 ms bins) with activity in V4 increasing shortly before IT, consistent with stimulus-related activity first occurring in V4. Figure 4B revisits our previous analyses (fig. 2C) but with spike counts binned into 10ms intervals rather than aggregated over the entire trial. Even with only 10ms of recordings, neural translation from V4 and IT to an advanced DCNN network layer minimises KL divergence between model outputs arising from image input vs. when driven by brain activity.

**Figure 4:**
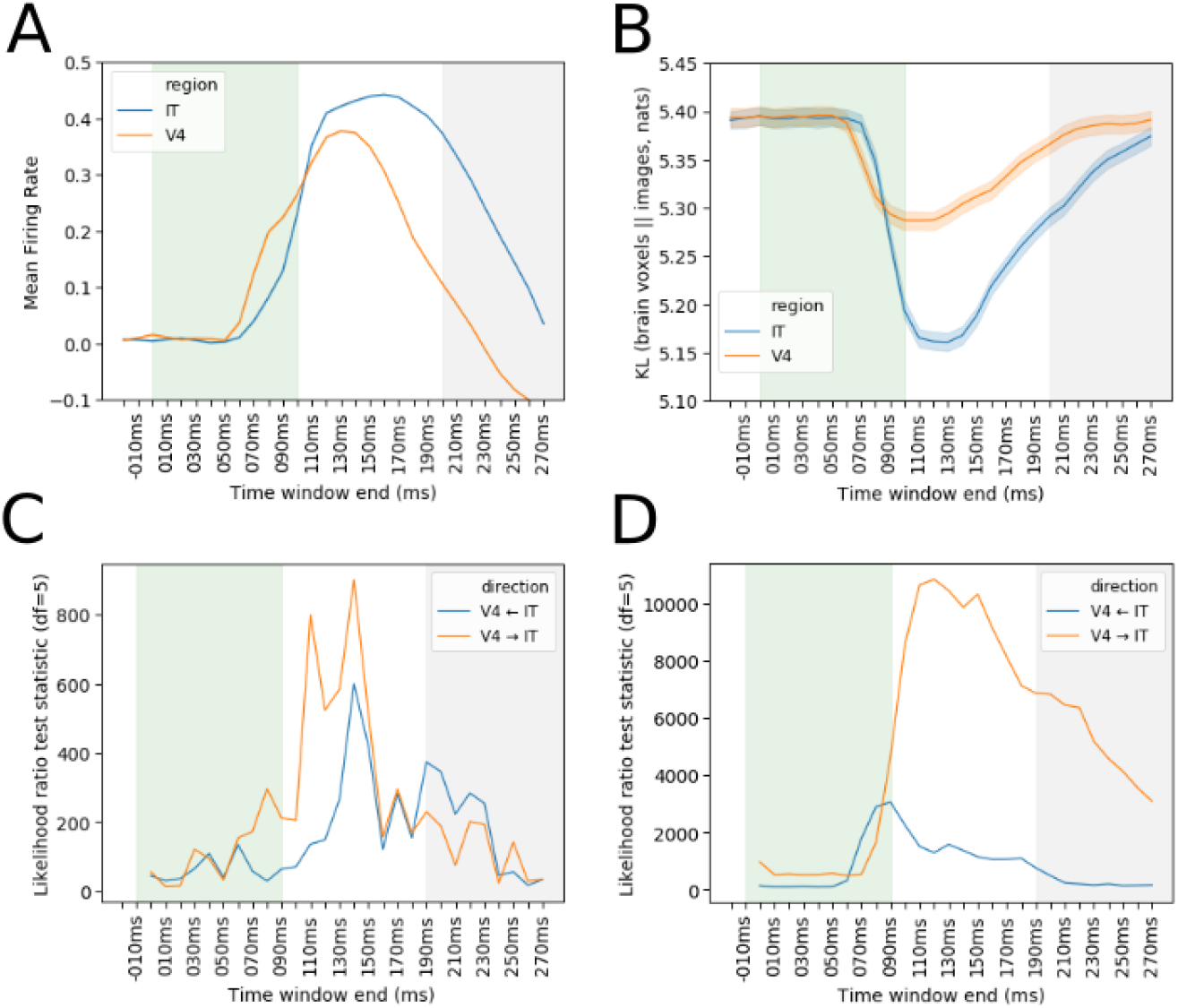
Analyses of monkey multi-unit recordings (*16*) time locked to stimulus presentation in 10ms time bins. Each visual stimulus was presented for 100ms (shaded green) with 100ms before the next (shaded grey). (**A**) Mean normalized spike counts for all electrodes for V4 and IT. (**B**) Task-relevant analysis (lower values imply closer correspondence with a late DCNN layer) show both V4 and IT can appropriately drive DCNN response (fig. 1B), starting around 70ms after stimulus onset. (**C**) Consistent with our long-range recurrence hypothesis (fig 3), Granger Causal Modelling indicates that, while V4 first drives IT in terms of raw firing rates (*V*4 → *IT*), (**D**) IT first drives V4 in terms of task-relevant information (*V*4 ← *IT*). These results are consistent with information about object category information (as assessed by interfacing with a late layer in a DCNN) first arising in IT and then feeding back to V4. At later time steps, Granger causality between V4 and IT becomes reciprocal (*V*4 ↔ *IT*) as the loop cycles.

Turning to the key Granger causality analyses, we evaluated whether early ventral stream regions become more like late-ventral stream regions over time due to recurrence (fig. 3). As processing unfolded, we found mutual causality between lower-level (V4) and higher-level (IT) areas for analyses conducted over spike counts (fig. 4C) and for analyses on the KL divergence times series that assessed the ability of brain regions to drive DCNN response (fig. 4D).

Critically, the specific predictions of the long-range recurrence hypothesis were supported with V4 first driving IT (*V*4 → *IT*) for the analysis of spike counts but IT first driving V4 (*V*4 ← *IT*) for the task-relevant information analysis using the KL divergence time series (see SI for details). These results are consistent with stimulus-driven bottom-up activity proceeding from V4 to IT on an initial feed forward pass through the ventral stream with actionable information about object recognition first arising in IT. Then, recurrent connections from IT to V4 make task-relevant information available to V4. As this loop is completed and cycles, both areas mutually influence one another with the impact of bottom-up stimulus information maintained throughout the process.

Computational models can help infer the function of brain regions by linking model and brain activity. Mulitlayer models, such as DCNNs, are particularly promising in this regard because their layers can be systematically mapped to brain regions. Indeed, the deep learning revolution in neuroscience began with analyses suggesting an early-to-early, late-to-late pattern of correspondence between DCNN layers and brain regions along the ventral visual stream during object recognition tasks (*3–5*).

However, as we have argued, correspondences based on total shared variance should be treated with caution. To complement these approaches, we presented a test focused on task-relevant variance that directly interfaced neural recordings with a DCNN model. If a brain region corresponds functionally to a model layer, then brain activity substituted for model activity at that layer should drive the model to the same output as when an image stimulus is presented. Of course, models and brains speak different languages, so a translation between brain and model activity must first be learned, which in our case was accomplished by a linear transformation. Once the translation function is learned, novel brain data and images can be used to evaluate possible brain-model correspondences.

Our approach, which focuses on task relevant variance within the overall computation, as opposed to local shared variance (fig. 1) uncovered a pattern of correspondences that dramatically differed from the existing literature. We found that all brain regions, from the earliest to the latest of visual areas along the ventral stream, best corresponded to later model layers. These results indicate that neural recordings in all regions contain higher-level information about object category even when most variance in a region is attributable to lower-level stimulus properties (fig. 3).

To resolve this discrepancy between our analyses focused on task-relevant variance and those based on shared variance, we evaluated the hypothesis that long-range recurrence between higher-level brain regions, such as IT, influenced activity in lower-level areas like V4. Analysing both firing rates of cells and information-level analyses using our brain-model interface approach, we found evidence that recurrent activity renders all areas functionally “late” as processing unfolds, even when total variance in some early visual regions is largely driven by bottom-up stimulus information. In this way, we integrate previous findings with our own and highlight how our method can be used to test hypotheses about information flow in the brain.

Our approach, which considers task-relevant variance, may help resolve conflicting interpretations on the function of brain regions. For example, the fusiform face area (FFA) responds selectively for faces, but its wider functional role in object recognition has been the subject of extensive debate (*17*). Here, we show that interfacing FFA into late model layers drives object recognition comparably to the lateral occipital complex (fig. 2B) on non-face natural images. We suspect that the function of a region will only be fully understood by considering task-relevant variance across several tasks in light of activity in connected brain regions. The tight interface we champion between computational models and brain activity should prove useful in evaluating theoretical accounts of how the brain solves tasks over time.

Computational models that perform the tasks end-to-end, from stimulus to behaviour, should be particularly useful. In essence, translating between brain regions to layers of such models can make clear what role a brain region plays within the overall computation. In the case of object recognition, our results suggested that recurrent models may be best positioned to explain how the nature of information within brain regions changes as the computation unfolds.

This conclusion is in line with a growing body of modelling work in neuroscience that affirms the value of recurrent computation (*18–20*). Unlike the aforementioned work, we suggest that long-distance recurrent connections that link disparate layers should be considered (cf. 21). We suspect such models will be necessary to capture time course data and the duality found in some brain regions, namely how most variance in a brain region can be attributable to lower-level stimulus properties while co-mingled with important higher-level, task-relevant signals.

As deep learning accounts in neuroscience are extended to other domains, such as audition (*22*), and language processing (*23*), the lessons learned here may apply. Our brain-model interface approach can help evaluate whether the brain processes signals across domains in an analogous fashion. By minding the distinction between shared and task-relevant variance, the role brain regions play within the overall computation may more readily come into focus.

Our approach may also have practical application in brain machine interfaces (BMI). Recent BMI developments have emphasised the readout of motor commands, neural processes taking place close to the periphery. In contrast, by leveraging the constraints provided by a pre-trained DCNN, we were able to gain traction on the ‘stuff of thought’, categorical and conceptual information in IT. Because we learned a general translation from brain to model, our approach applied to BMI would allow distant generalisation. For example, we were able to extrapolate to novel categories (fig. S3). For example, a translation from brain to model that never trained on horses, but trained on other categories, can perform zero-shot generalisation when given brain activity elicited by an image of a horse. The interface has the potential to produce a domain-general mapping rather than one dependent on specific training data. In the future, BMI approaches that address general thought without exhaustive training on all key elements and their combinations may be feasible.

## Acknowledgments

The authors thank colleagues in the LoveLab for discussion and comments on early versions of this manuscript.

## Author Contributions

N.J.S: Conceptualization, methodology, software, validation, formal analysis, investigation, data curation, writing - original draft, writing - review & editing, visualization. B.C.L.: Conceptualization, methodology, resources, writing - review & editing, supervision, funding acquisition.

## Supplementary materials

Materials and Methods

Figs. S1 to S4

Tables S1 to S2

References (*26-33*)

