## Supplementary Information for "Directly interfacing brain and deep networks exposes non-hierarchical visual processing"

### S1 Materials and Methods

#### Datasets

We re-analysed three existing neural datasets. Two, BOLD5000(*14*) and Generic Object Decoding(*15*) consist of fMRI from human subjects who viewed images taken from Imagenet(*24*) a benchmark large dataset of natural images. We restricted the BOLD5000 dataset to only those images drawn from Imagenet (2012 ILSVRC) edition and to subject 1-3 who completed the full experiment. The analysis of Generic Object Decoding used the data from the 'training' portion of their image presentation experiment, consisting of 1200 images from 150 categories drawn from the Imagenet Fall 2011 edition. For both datasets, each image was presented once, thus each row represents individual trials.

The third dataset consists of neuron spike counts directly recorded from V4 and IT of two macaque monkeys (*16*) in a rapid serial visual presentation paradigm where each image is passively viewed for 100ms, with 100ms between images. We used the publicly available data processed as detailed in those publications. For the neural interfacing analysis of the spiking neural dataset, we used spike rates aggregated over multiple presentations of each of 3200 unique images, in the interval 70-170ms after stimulus onset, with the electrodes from the two subjects concatenated, as in the original analysis (*16*) For the Granger causal modelling analysis of the same dataset, we used spike rates at the level of the individual trial (i.e., no aggregation) for each 10ms time bin.

The neural data corresponding with each image was related to layer activations of a deep convolutional neural network (DCNN) trained on image classification, when processing the same pixel-level data. The three neural datasets contain data for various brain regions from ventral stream, including visual areas (V1, V2, V3 V4, included as 'EarlyVis' in *14*), areas responsible for processing shape and conceptual information (LOC, IT) and various downstream areas (OPA, PPA, FFA, RSC).

For details on neuroanatomical placement or functional localisation of each region, we refer readers to the original publications. Further details of brain regions and dimensionality of the data from each region are presented in supplementary information table S1. 1.

#### DCNN

As the base DCNN for all simulations, we used a re-implemented and trained version of VGG-16 (*13*, configuration D) using Keras(*25*) version 2.2.4 and TensorFlow version 1.12. This model was selected for its uncomplicated architecture, near-human level classification accuracy on ImageNet, and widely reported robust correspondence with primate or human data on various measures, including human behavioural (similarity judgements (*26*) human image matching (*20*), and neural (*20*, *27*). We implemented and trained a version of the architecture with $64 \times 64 \times 3$ input size, with corresponding changes in spatial dimensions for all layers (table 2). For all analyses, images from all datasets were cropped to a square and resized to this resolution.

For the monkey multi-unit dataset, where images are contained in a circular frame, the central $192 \times 192$ portion of the $256 \times 256$ original was cropped and resized, to decrease the proportion of image taken up by blank space in the corners. While the original authors trained their network in a two-stage process, beginning with a subset of the layers, the inclusion of batchnorm(*28*) between the convolution operation and activation function of each layer enabled training the complete network in a single pass. We used the authors setting for weight decay ($\ell_2$ penalty coefficient of $5 \times 10^{-4}$) and a slightly different value for dropout probability (0.4). Model architecture details are presented in the supplementary information (table S2).

#### DCNN training

Our training procedure followed (*13*). The model was trained on ImageNet 2012 (1000 classes) for analyses of the BOLD5000 and monkey multi-unit datasets. For the Generic Object Decoding dataset, the model was trained to convergence on ImageNet Fall 2011 (21841 classes), before layer FC3 was replaced and retrained with 150 classes, corresponding with the classes used in our re-analysis of (*15*) For ImageNet Fall 2011 we randomly allocated 2% of each class including all images used in (*15*) to an in-house validation set that was not used for training. One image used in the original study was missing from our image dataset and was excluded from all analyses. All images were resampled from their native resolution to $64 \times 64 \times 3$ by rescaling the shortest side of the image to 64 pixels and centre-cropping.

Both versions of the model was trained using mini-batch stochastic gradient descent, with a batch size of 64, an initial learning rate of 0.001 and Nesterov momentum of 0.90561. The learning rate decayed by a factor of 0.5 when validation loss did not improve for 4 epochs, with training terminating after 10 epochs of no improvement. All layers used Glorot normal initialization. During training, images were augmented with random rescaling, horizontal flips and translations.

#### Cross-validation

Classifier-based methods require training classifier parameters, before evaluating it on data withheld from the training set. In all analyses, we use the standard approach of k-fold cross validation(*29*) in which the dataset is randomly allocated into $k$ equally-sized partitions, and the analysis is iterated $k$ times, each time training on $k-1$ partitions and evaluating on one. In this way, the classifier is evaluated over the entire dataset. For all analyses, except where otherwise specified, we use stratified 8-fold cross validation, that is to say dataset items are randomly allocated to partitions with the constraint that $1/k$ of each class be allocated to each validation partition. For the spiking neural dataset(*16*), each unique image was rendered from one of 64 objects, with varying position and orientation. Here, stratification was done at the object level.

For the out-of-training-class generalisation analysis, we used leave-one-class-out cross validation, where for $m$ classes, the analysis is iterated $m$ times, the evaluation set consisting only of the entirety of a single class, on each iteration.

#### Neural Interfacing analysis

Given a dataset $D$, consisting of an image matrix $D_i$ of shape $(n, 64, 64, 3)$ where n is the number of images, and a corresponding neural data matrix $D_r$, of shape $(n, d)$ where $d$ is the number of neural features (electrodes, for multi-unit data, or voxels, for fMRI data), consider a DCNN computing a function $f$ on $D_i$, mapping $D$ to $P_i$, an $(n, m)$ matrix of predictions, each row being a probability distribution over the $m$ classes the DCNN was originally trained to classify.

$$f(D_i) = P_i \tag{1}$$

For an arbitrary intermediate model layer $q$, we may decompose $f$ into $g_q$ and $g'_q$, by computing intermediate activations, $g_q(D_i)$:

$$f(D_i) \equiv g'_q(g_q(D_i)) = P_i \tag{2}$$

The neural interface analyses proceeded by applying a linear transform $W$ to the centered and column-normalized neural data, $D_r$ and inputting the result into DCNN layer $q$, to compute a matrix of model predictions for the neural data, $P_r$.

$$g'_q(WD_r) = P_r \tag{3}$$

##### Linear transformation matrix training

The transformation matrix $W$ was computed by partitioning image and neural datasets $D_i, D_r$ into training and evaluation partitions using 8-fold cross-validation, and $W$ was learned as a linear mapping from $D_r$ to the layer $q$ activations generated by the corresponding images, $D_i$, on the training partition:

$$g_q(D_i) = WD_r + \epsilon \tag{4}$$

For each cross-validation fold, the model predictions were computed for the evaluation partition. In practice, $W$ was computed as a single-layer linear neural network with no bias or activation function, to minimise mean-squared error of supervision targets $g_q(D_i)$ using mini-batch stochastic gradient descent with momentum, (batch size 64, momentum of 0.9, $l_2$ regularization of 0.0003, initial learning rate of 0.1, decreasing by a factor of 0.5 when validation loss did not improve for 4 epochs and terminating after 400 epochs or after validation loss did not improve for 20 epochs.) For the analysis of the macaque dataset(*16*), on the level of the individual trial, prior to performing the GCM model, W was computed using the Adadelta optimizer (batch size of 128, initial learning rate of 0.04.)

We also considered an alternative mode for training $W$, by first assembling the model in the form of equation 3, composed of transformation matrix $W$ initialised with small random weights, followed by DCNN layer $q$ onwards, $g'_q$, thus mapping end-to-end from neural measures $D_r$ to output. $W$ was then trained by back-propagating the categorical cross-entropy error term from the softmax output layer, using the supervision target of the ground-truth labels for the neural dataset ($D_r$), with all other weights in the network frozen. This method produced a pattern of results that was qualitatively similar, although with lower absolute accuracy (fig. S8).

##### Neural Interface Evaluation

The output of the model, $P$, is an $(n, m)$ matrix of probability distributions over the $m$ output classes the original DCNN was trained on, for each of $n$ images in $D$. We computed this for the original DCNN on the image dataset, $f(D_i) = P_i$, and also for the neural dataset for each brain region $r$ and model layer $q$, $g'_q(WD_r) = P_r$. The correspondence between $r$ and $q$ was evaluated by comparing the model predictions $P_r$ either against the ground-truth classes (by computing the overall AUC of the classifier, via the equality between AUC and Wilcoxon-Mann-Whitney $U$) or against model predictions from the image dataset, by computing the KL divergence of $P_r$ from $P_i$ for each row $n$.

#### Shared Neural Variance Analysis

For comparison, we present an example of a shared neural variance analysis using the macaque spiking neuron dataset(*16*) and our re-implemented model. Conceptually, in common with the interfacing analysis (section ), the analysis evaluates the correspondence between a brain region $r$ and a model layer $q$. Layer $q$ model activations, $g_q(D_i)$, were compared with a neural dataset obtained from the presentation of corresponding images, $D_r$. To establish our results are comparable to those previous, we used the neural predictivity method exactly as implemented in the Brainscore benchmark for DCNNs (*30*).

The dataset was iteratively partitioned using 8-fold cross-validation into training/validation partitions. Following the method of (*23*), we used the image stimuli from the training partition to generate model activations on each layer. We used PCA to calculate the first 1000 principal components of these activations, before training a PLS regression model (25 components) to predict, for each electrode, the firing rate across the validation partition. The predictivity for each electrode was computed as the Pearson correlation coefficient between the predicted firing rates across the dataset and the actual recorded values, with the overall predictivity given by the correlation coefficient of the median electrode.

#### Simple Classifiers on the Neural Datasets

To establish performance baselines for the interfaced fMRI datasets, which were evaluated in terms of classification performance, we applied various standard classifiers to the neural data directly, to predict the image class from the neural data from various brain regions. Known as multi-voxel pattern analysis (MVPA), evaluating the trained classifier's ability to predict class labels from fMRI or spiking neural data is now a standard approach to quantifying the categorical-level information within a brain region(*31*). Nevertheless, in the present analyses the number of different classes is unusually large, and the number of examples from each class unusually small, (1916 images from 958 classes(*14*) 1200 images from 150 classes(*15*) for a straightforward MVPA analysis on these datasets. We report the AUC of the classifier computed in the same way as for the neural interfacing analysis . All classifiers were implemented as detailed below using version 0.20.3 of the Scikit-Learn library(*32*).

##### Multiclass Logistic Regression

Implemented as LogisticRegression with the 'multinomial' option, the lbfgs solver and a maximum of $10^3$ iterations.

##### Nearest Neighbours Classifier

Implemented as KNeighboursClassifier. Given the structure of the BOLD5000 dataset, with only two examples per class (thus, either one or two examples in the training partition, test classification of each class on the basis of one correct training example) we classified on the basis of the single nearest neighbour under a Euclidean distance function.

##### Linear Support Vector Machine (SVM)

Implemented as LinearSVC, using a one-versus-rest multi-class strategy, with a maximum of $10^4$ iterations and $C$ parameter of $10^{-3}$.

#### Granger Causal Modelling

In contrast to the previous neural interfacing analysis of the spiking neural dataset, which aggregated spike rates over multiple presentations of each image, in the interval 70-170ms after stimulus onset, here we trained and evaluated the model on data at the individual trial level. We conducted a separate decoding analysis for each 10ms time bin, from -20ms (i.e., prior to stimulus onset) to 270ms, with all time indices referring to the preceding 10ms bin. Training linear transformation matrix $W$ is described in section . Prior to the GCM, we pre-processed the trial-level relative entropy data to ensure stationarity by, first, subtracting the temporal mean and standard deviation from each trial, and second, subtracting the mean signal and dividing by the signal's standard deviation, thus ensuring that each time step has zero mean and unit variance.

Given two regions, X and Y, separate Granger-Causal models were computed for each direction $X \rightarrow Y$ and $X \leftarrow Y$, where each model takes the form of a linear regression, where the univariate outcome:

$$KL(D_X||D_i)_n \tag{5}$$

the KL divergence of region X with $\theta$, the base model predictions, is predicted by the Granger null model (6), or the Granger-causal model (7).

$$KL(D_X||D_i)_{n-1}, ..., KL(D_X||D_i)_{n-p} \tag{6}$$

$$KL(D_X||D_i)_{n-1}, KL(D_Y||D_i)_{n-1}, ..., KL(D_X||D_i)_{n-p}, KL(D_Y||D_i)_{n-p} \tag{7}$$

where $p$, the maximum number of previous time-steps is a hyperparameter that is determined using model-selection criteria such as BIC. The appropriate model was determined by comparing log-likelihood ratios, given the data, for the causal and null models.

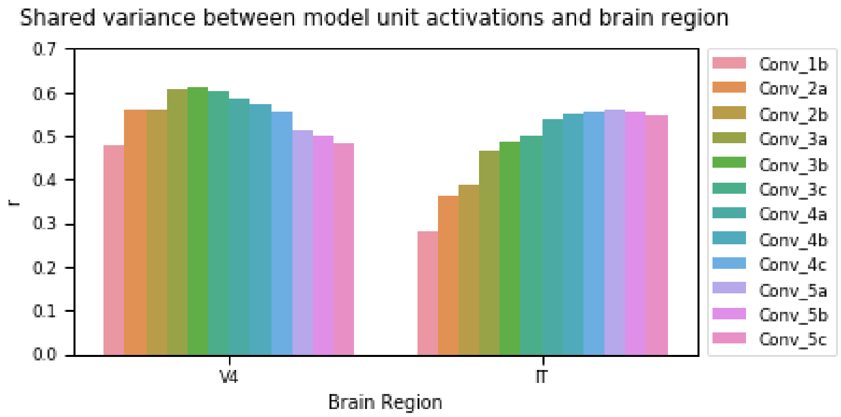


Figure 1: **Fig. S1:** Standard approaches to relating primate ventral stream and DCNNs evaluate variance shared between data from each brain region and unit activations on each DCNN layer. They have been taken as evidence that earlier ventral stream regions (e.g., V4) correspond to earlier DCNN layers and later regions (e.g., IT) correspond to later DCNN layers. Here, we present a shared-variance based analyses of directly recorded spiking neural activity (*16*) and VGG-16 using an established method (*20*) Higher correlations reflect more shared variance between brain region and model layer.

### SI Figures

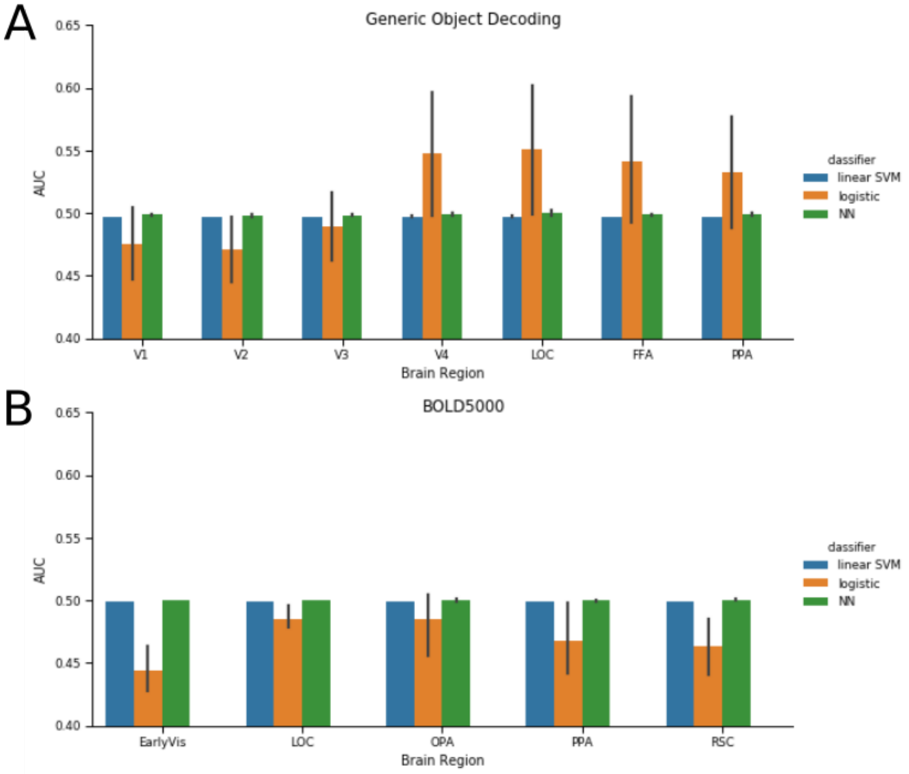


Figure 2: **Fig. S2:** Comparison of the brain-DCNN interface as a neural pattern classifier, compared with standard linear classifiers typically used in multi-voxel pattern analysis (MVPA). We present classification performance (AUC) directly on neural patterns, on the Generic Object Decoding (**A**) and BOLD5000 (**B**) datasets, for simple classifiers (support vector machine with linear kernel, multiclass logistic regression, and a 1-nearest neighbour classifier) with results for interface with each layer of the DCNN presented for comparison. Performance of the simple classifiers is generally near chance (0.5), we attribute this to the large number of image classes (150, 958 respectively) and few available examples (2, 8 per class) which severely limit the available training data. Because the brain-DCNN interface learns a general mapping between brain region and model, it does not suffer this limitation, making it an appealing novel approach for MVPA.

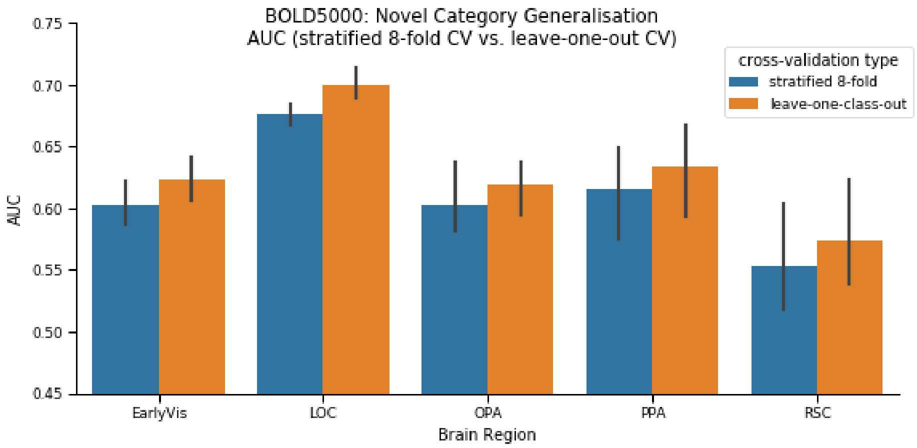


Figure 3: **Fig. S3:** Learning a mapping directly from neural measures to DCNN activation space produces a general mapping, rather than being dependent on training examples. The neural interface has an in-built ability to generalise to novel classes. This is demonstrated by presenting classification performance (AUC) on the BOLD5000 dataset, by comparing cross-validation (CV) strategies. Error bars represent 95% confidence intervals across 3 subjects. Stratified 8-fold CV (the default, used in all other analysis) ensures each training partition contains at least one example of each class. Leave-one-class-out CV involves the same number of CV folds as there are classes, each time training on all data except one class, which is withheld for the validation set. Performance is equivalent or better (LOC) when generalising to novel classes, which we attribute to more training data per CV fold. Due to the training time, this analysis was restricted to layer 5a.

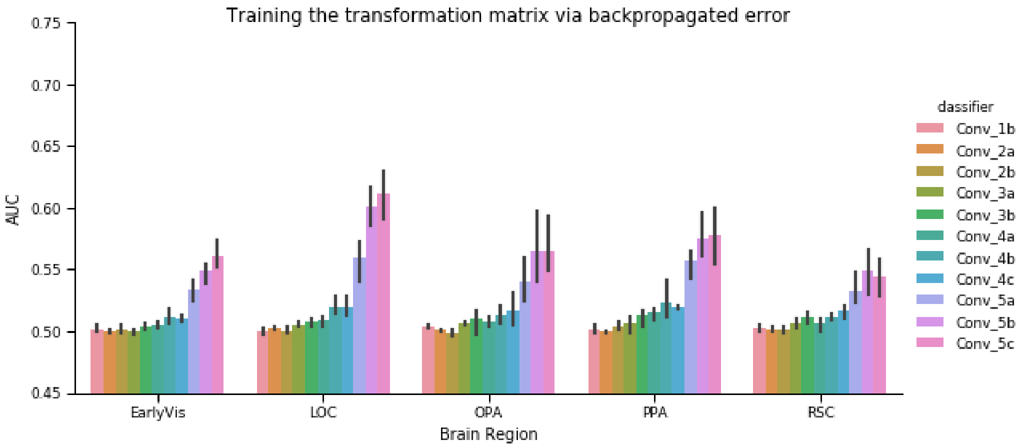


Figure 4: **Fig. S4:** Alternative 'backprop mode' for training the transformation matrix $W$ mapping from neural space to DCNN activation space. Classification performance (AUC) on the BOLD5000 dataset follows a qualitatively similar pattern to the main analysis (compare fig. 2B), albeit with lower absolute accuracy. The default analysis trains $W$ independently as a regression problem, using layer activations as supervision targets directly. Instead, this approach uses $W$ as a weights matrix for a new neural network that takes neural data from a brain region as input, connected to the latter part of the DCNN, and training the network using the class labels as supervision targets, with all other DCNN weights frozen.

### SI Tables

| Dataset | Generic Object Decoding(*15*) | BOLD5000(*14*) | Linear Weighted Sums(*16*) |
|---|---|---|---|
| Stimuli | experiment 'train' phase: 1200 images from 150 categories (ImageNet Fall 2011) | 1916 images from 958 categories (ImageNet ILSVRC 2012 | 3200 greyscale composite images, 64 objects in 8 categories,) non-congruent background |
| Task | one-back repetition detection | valence judgement ('like', 'neutral', 'dislike') | passive viewing, RSVP presentation 100ms/100ms |
| Subjects | 5 human fMRI | 3 human fMRI (partial data from subject 4 excluded) | 2 Macaque monkeys (vectors concatenated) multi-unit recording |
| Time indices | full 9s of image presentation | TR3-4 | 70-170ms |
| Brain region (dimensionality per subject) | V1 (1004, 757, 872, 719, 659)<br>V2 (1018, 944, 1031, 855, 891)<br>V3 (759, 810, 861, 929, 907)<br>V4 (740, 544, 754, 704, 860)<br>LOC (540, 834, 996, 668, 566)<br>PPA (356, 316, 496, 398, 550)<br>FFA (568, 435, 928, 725, 929) | EarlyVis (495, 495, 1218)<br>LOC (342, 888, 1027)<br>OPA (288, 180, 392)<br>PPA (331, 370, 273)<br>RSC (229, 421, 394) | IT (168 = 58 + 110)<br>V4 (88 = 70 + 18) |

Table 1: **Neural datasets** For further dataset details, such as how regions were defined, we refer readers to the original publications

| Block | Layer | Dimensions ($h \times w \times c$) | Filter Size |
|---|---|---|---|
| Input | | $64 \times 64 \times 3$ | |
| 1 | 1a | $64 \times 64 \times 64$ | $3 \times 3$ |
| | 1b | $64 \times 64 \times 64$ | $3 \times 3$ |
| | max pool 1 | | $2 \times 2$ |
| 2 | 2a | $32 \times 32 \times 128$ | $3 \times 3$ |
| | 2b | $32 \times 32 \times 128$ | $3 \times 3$ |
| | max pool 2 | | $2 \times 2$ |
| 3 | 3a | $16 \times 16 \times 256$ | $3 \times 3$ |
| | 3b | $16 \times 16 \times 256$ | $3 \times 3$ |
| | 3c | $16 \times 16 \times 256$ | $3 \times 3$ |
| | max pool 3 | | $2 \times 2$ |
| 4 | 4a | $8 \times 8 \times 512$ | $3 \times 3$ |
| | 4b | $8 \times 8 \times 512$ | $3 \times 3$ |
| | 4c | $8 \times 8 \times 512$ | $3 \times 3$ |
| | max pool 4 | | $2 \times 2$ |
| 5 | 5a | $4 \times 4 \times 512$ | $3 \times 3$ |
| | 5b | $4 \times 4 \times 512$ | $3 \times 3$ |
| | 5c | $4 \times 4 \times 512$ | $3 \times 3$ |
| | max pool 5 | | $2 \times 2$ |
| FC | FC1 | 4096 | |
| | dropout 1 | | |
| | FC2 | 4096 | |
| | dropout 2 | | |
| | FC3 (output) | 1000 softmax | |

Table 2: **DCNN Architecture:** Layer configuration and dimensions of the DCNN used for all analyses.
